# Potently neutralizing human monoclonal antibodies against the zoonotic pararubulavirus Sosuga virus

**DOI:** 10.1101/2022.11.17.516973

**Authors:** Helen M. Parrington, Nurgun Kose, Erica Armstrong, Laura Handal, Summer Diaz, Joseph Reidy, Jinhui Dong, Guillaume B. E. Stewart-Jones, Punya Shrivastava-Ranjan, Shilpi Jain, César G. Albariño, Robert H. Carnahan, James E. Crowe

**Author notes:** **Address correspondence to:** James E. Crowe, Jr., Vanderbilt Vaccine Center, 11475 Medical Research Building IV, 2213 Garland Avenue, Nashville, TN 37232-0417, USA.

## Abstract

Sosuga virus (SOSV) is a recently discovered paramyxovirus with a single known human case of disease. There has been little laboratory research on SOSV pathogenesis or immunity, and no approved therapeutics or vaccines are available. Here, we report the discovery of human monoclonal antibodies (mAbs) from the circulating memory B cells of the only known human case and survivor of SOSV infection. We isolated six mAbs recognizing the functional attachment protein (HN) and 18 mAbs against the fusion (F) protein. The anti-HN mAbs all target the globular head of the HN protein and can be organized into 4 competition-binding groups that exhibit epitope diversity. The anti-F mAbs can be divided into pre- or post-fusion conformation-specific categories and further into 8 competition-binding groups. Generally, pre-fusion conformation-specific anti-F mAbs showed higher potency in neutralization assays than did mAbs only recognizing the post-fusion conformation of F protein. Most of the anti-HN mAbs were more potently neutralizing than the anti-F mAbs, with mAbs in one of the HN competition-binding groups possessing ultra-potent (<1 ng/mL) half maximal inhibitory (IC_50_) virus neutralization values. These findings provide insight into the molecular basis for human antibody recognition of paramyxovirus surface proteins and the mechanisms of SOSV neutralization.

## Introduction

Paramyxoviruses are enveloped, single-stranded, negative-sense RNA viruses with a broad range of hosts including mammals, birds, and fish (1). Paramyxoviruses have a long evolutionary history replicating in various bat species (2), and bats have been the source of several zoonotic paramyxovirus spillovers into human populations (3–6). Therefore, understanding human immune responses against zoonotic and endemic paramyxoviruses is important for increasing our knowledge of the fundamental basis of immunity to these viruses and for informing development of candidate therapeutic antibodies or vaccines.

Sosuga virus (SOSV) is a recently discovered zoonotic paramyxovirus that caused a near-fatal, acute febrile disease in a wildlife researcher who had been conducting surveillance studies in South Sudan and Uganda (7, 8). The source of the infection was traced back to Egyptian rousette (*Rousettus aegyptiacus*) bats, which likely serve as an animal reservoir of this virus (8). As SOSV only recently emerged, much remains unknown about this virus including its potential to cause human epidemics. The severe disease in the index case and the high mortality rates of other bat-borne paramyxoviruses, such as Nipah virus (9, 10) suggest that SOSV may have the potential to cause epidemics of life-threatening disease.

SOSV is classified as a paramyxovirus in the *Rubulavirinae* subfamily, and the viral genome encodes the hemagglutinin-neuraminidase (HN) and fusion (F) glycoproteins (8, 11, 12). Both surface proteins are necessary for viral entry into host cells. The HN protein recognizes and binds the viral receptor, and the F protein mediates viral fusion with host-cell membranes (11, 12). While the HN proteins of orthorubulaviruses, such as mumps virus, are known to bind to sialic acid (11, 13, 14), this is not true of pararubulaviruses, including SOSV, which do not bind sialic acid (15, 16). To date, the receptor of SOSV and other pararubulaviruses remains unknown. As the two glycoproteins are expressed on the surface of the viral envelope, they also are vulnerable as targets for recognition by neutralizing monoclonal antibodies (mAbs). The neutralizing antibody response to related paramyxoviruses is predominantly targeted to HN (17–21), but the most commonly targeted antigen recognized following natural SOSV infection and the mechanism by which SOSV-neutralizing antibodies function are unknown. It is possible that structurally conserved antigenic sites exist across diverse species of rubulaviruses due to structural similarities between their F and HN proteins and the essential role of these proteins in viral fusion (11, 12).

Here, we sought to investigate the human antibody response against SOSV surface glycoproteins and identify antibody-mediated mechanisms of neutralization and breadth of reactivity of mAbs from the only known human survivor of SOSV infection. We are not aware of any previous reports of isolation of human mAbs against a rubulavirus. Such studies may contribute to a deeper understanding of the human immunity to paramyxoviruses and could have direct applications in preventing or diagnosing outbreaks of SOSV. We isolated potently neutralizing mAbs from circulating memory B cells and determined the epitope specificity, binding affinity, neutralization potency and cross-reactivity of the mAbs between diverse rubulaviruses. The data reveal key information on the neutralizing immune response to a novel paramyxovirus and identify pan-rubulavirus epitopes recognized by some human mAbs. The mAbs generated in this study also could be used as candidate prophylactic or therapeutic antibodies, diagnostic reagents, or as research tools for future studies on SOSV and related viruses.

## Results

### Cell-surface expression of recombinant HN and F SOSV protein antigens

Wild-type protein-encoding sequences for the two major surface proteins of SOSV (F and HN) were synthesized as cDNA and cloned into a mammalian expression vector (pTwist-CMV) by Twist. These constructs were transfected singly or in combination into Vero cells, and by 48 hr visible syncytia were observed in the wells co-transfected with F and HN but not in un-transfected cells or wells containing cells transfected with only HN-encoding or only F-encoding plasmids (**Figure 1**). Syncytium formation mediated by HN + F expression suggested both proteins were expressed on the cell surface and in a functional state.

**Figure 1.**
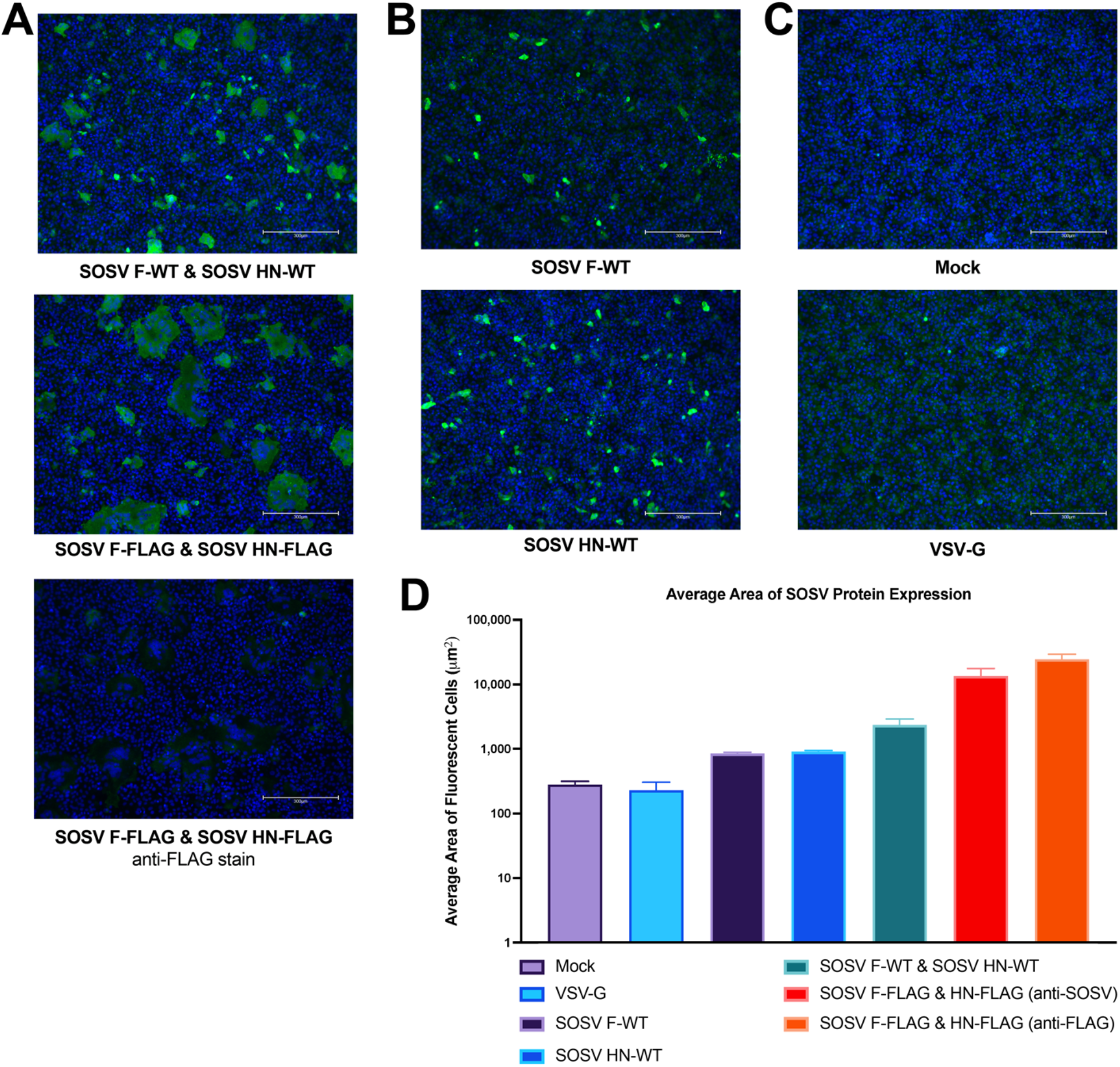
Co-transfection of cDNAs encoding SOSV F and HN proteins causes robust syncytia formation in cell culture monolayers. Representative field of view (10× objective) of transfected Vero cell culture monolayers. Nuclei were stained with 4’,6-diamidino-2-phenylindole (DAPI; blue) and SOSV proteins were stained with a polyclonal mix of six anti-SOSV mAbs (three anti-HN and three anti-F) or mouse anti-FLAG antibody with goat anti-human IgG with Alexa Fluor 488 dye or goat anti-mouse IgG with Alexa Fluor 488 dye antibodies as secondary antibodies. **(C)** Syncytia producing transfections: Co-transfection of SOSV F-WT + SOSV HN-WT, co-transfection of SOSV F-FLAG + SOSV HN-FLAG, or co-transfection of SOSV F-FLAG + SOSV HN-Flag constructs stained with anti-FLAG antibodies. **(B)** Non-syncytia producing transfections: cDNA encoding SOSV F-WT or SOSV HN-WT were transfected individually. **(C)** Controls: mock transfection or VSV G-WT transfection. **(D)** Average area of fluorescently stained clusters (cells or syncytia).

### Expression of recombinant HN SOSV antigens for production of soluble proteins

We developed expression systems for soluble SOSV HN proteins by truncating the wild-type HN sequence to encode amino acid residues 75 to 582 comprising the HN protein ectodomain (designated HN_ecto_) or residues 125 to 582 comprising the HN protein head domain (designated HN_head_) (**Figure 2**), adding a CD5-signal peptide sequence and thrombin-cleavable 6× His tag to the amino terminus of the proteins. These constructs were sequence-optimized for human cell expression, synthesized as cDNA, and cloned into the pTwist-CMV mammalian expression vector (Twist Biosciences). Soluble HN proteins were expressed in Expi293F cells for 5 to 7 days and purified from cell supernatants using an ÄKTA pure system (Cytiva), and the eluate was concentrated and buffer-exchanged into Dulbecco’s Phosphate Buffered Saline (DPBS). As expected, the final purified proteins were about 70 kDa in apparent molecular weight.

**Figure 2.**
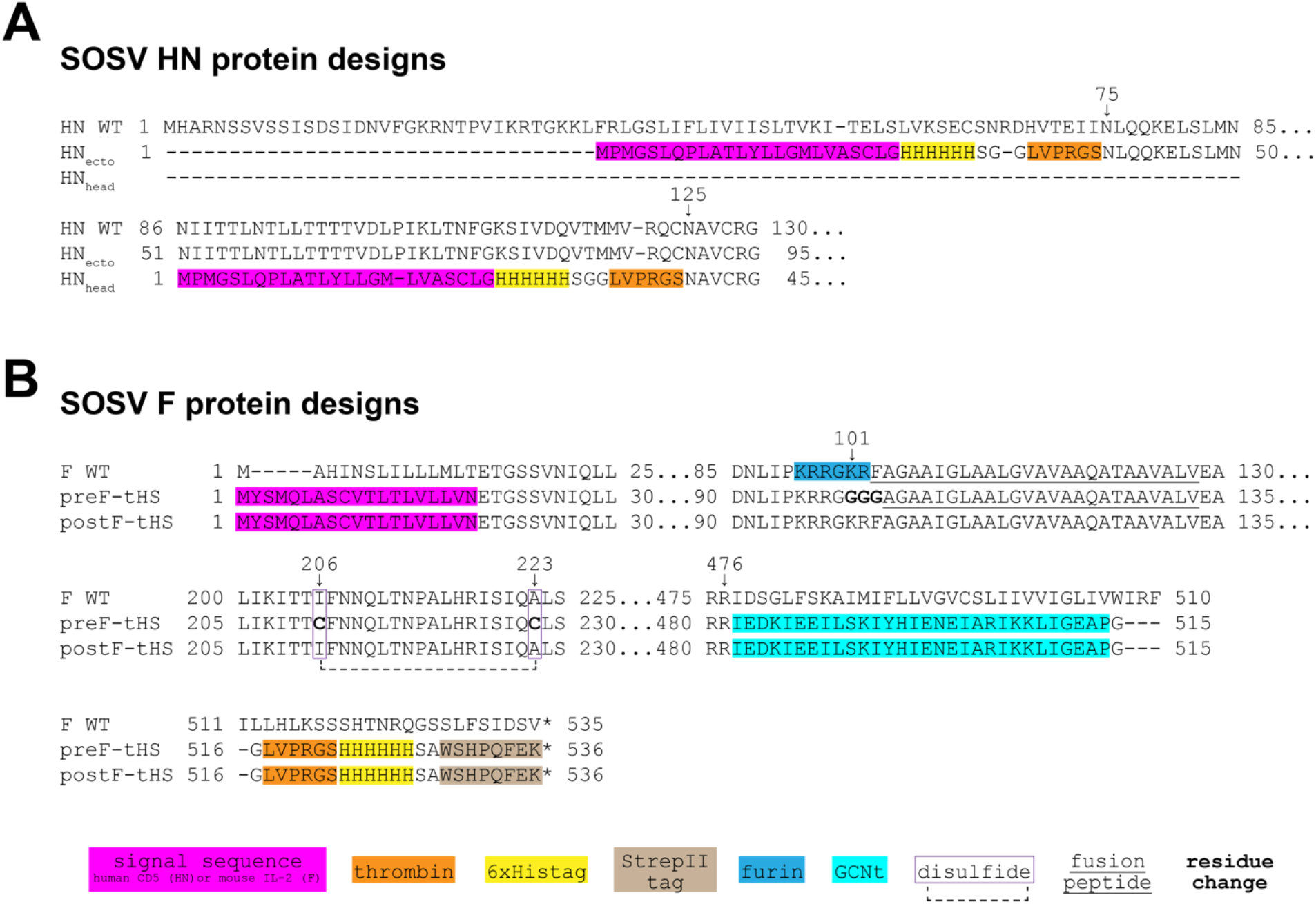
SOSV HN and F soluble protein designs. Soluble versions of the SOSV HN and F proteins were generated by removing the cytoplasmic tail and transmembrane domains and adding a human CD5 or mouse IL-2 signal peptide. To aid in purification, 6× His tag or StrepII tags were added to the carboxy (F) or amino (HN) terminal ends. **(A)** HN_ecto_ design includes a longer portion of the stalk region starting at residue 75, while the HN_head_ construct is comprised of almost only the globular head domain of the HN protein. **(B)** Additional modifications were necessary to include in the SOSV F pre-fusion (preF-tHS) construct which included removal of the furin cleavage site and creation of a stabilizing disulfide bond by point mutations to cysteines at 206 and 223. The post-fusion construct of SOSV F (postF-tHS) more closely resembles the wild-type sequence but with replacement of the cytoplasmic and transmembrane domains with a GCNt trimerization domain (also in preF-tHS).

### Expression of recombinant F SOSV antigens for production of soluble proteins

We designed soluble forms of the SOSV F protein in both pre-fusion-stabilized and post-fusion conformations and closed the corresponding synthesized cDNAs into the pVRC4800 vector. Both the pre- and post-fusion F protein constructs contain a mouse IL-2 signal sequence, a truncated the SOSV F protein sequence at residue 467, and an engineered, trimeric coiled-coil domain of the yeast transcriptional activator GCN4 leucine zipper (GCNt (22)) replacing the transmembrane and cytoplasmic domains. Additionally, the pre-fusion design disrupted the poly-basic furin cleavage site by replacing three residues (from 101-103) with a soluble glycine linker and adding a stabilizing disulfide bond between residues 206 and 223. Soluble protein was generated by expressing the constructs in Expi293F cells for 5 days before purification on ÄKTA pure system (Cytiva) or AZURA P 6.1L (Knauer).

### Isolation of SOSV-reactive mAbs

Whole blood samples from the only known human case of SOSV infection were obtained five years after infection following informed written consent. Peripheral blood mononuclear cells were isolated and transformed with Epstein-Barr virus to form lymphoblastoid cell lines, expanded, fused with a non-secreting myeloma cell partner to make stable hybridoma cell lines, and single cell-sorted using flow cytometry to isolate biologically cloned hybridoma cell lines secreting mAbs. Throughout this process, cell supernatants were screened for binding to F or HN cell-surface expressed SOSV glycoproteins to identify the cell lines containing SOSV-specific IgG-secreting B cells. In total, 24 SOSV-reactive mAbs were isolated, with 18 binding to F protein and 6 binding to HN protein (**Table 1**). The antibodies are predominantly of the IgG1 subclass, with two mAbs (SOSV-2 and −32) being IgG3. Most (19 of the 24) mAbs use a k light chain, while five clones (SOSV-13, −21, −64, −83, and −85) use a λ_1_ or λ_2_ light chain. Recombinant versions of each member of the mAb panel were created by synthesizing a cDNA encoding the antibody variable gene regions and cloning by Gibson assembly into a human IgG1 expression vector (with k or λ light chain as appropriate). The recombinant mAbs (rmAbs) each exhibited the ability to bind the appropriate cell-surface displayed viral antigen, as expected, and were used in all assays in this work.

**Table 1.** Half-maximal effective concentration (EC_50_) values for antibody binding to recombinant soluble F or HN proteins in ELISA.

| SOSV protein specificity | MAb (rSOSV- ) | EC<sub>50</sub> value for binding (ng/mL) to indicated protein antigen (± SD) |  |
| --- | --- | --- | --- |
| **F** |  | **Pre-fusion F** | **Post-fusion F** |
|  | 2 | 26.5 ± 9.4 | 42.7 ± 18.5 |
|  | 5 | 25.4 ± 23.0 | 1.7 ± 0.8 |
|  | 10 | 23.4 ± 14.9 | 1,936 ± 458 |
|  | 21 | 37.7 ± 11.5 | > |
|  | 23 | 19.4 ± 6.6 | 18.5 ± 2.0 |
|  | 32 | 42.0 ± 14.0 | 17.0 ± 3.5 |
|  | 35 | > | > |
|  | 38 | 19.3 ± 2.3 | 4,546 ± 2,687 |
|  | 39 | 28.3 ± 14.8 | > |
|  | 44 | 20.9 ± 6.6 | 1.0 ± 0.3 |
|  | 53 | 19.1 ± 3.9 | 1.8 ± 1.8 |
|  | 59 | > | > |
|  | 64 | 54.2 ± 20.1 | N/A |
|  | 66 | 31.8 ± 7.5 | N/A |
|  | 68 | 23.3 ± 15.3 | 1.5 ± 0.4 |
|  | 73 | 48.5 ± 9.2 | 951 ± 230 |
|  | 77 | 15.7 ± 5.3 | 1.0 ± 0.4 |
|  | 85 | 1,209.6 ± 1,184.0 | 12.6 ± 5.6 |
| **HN** |  | **HN<sub>ecto</sub> protein** | **HN<sub>head</sub> protein** |
|  | 13 | 38.9 ± 27 | 29.5 ± 8.7 |
|  | 19 | 28.1 ± 9.6 | 22.0 ± 5.7 |
|  | 24 | 66.4 ± 41 | 64.0 ± 26 |
|  | 29 | 78.8 ± 24 | 92.2 ± 21 |
|  | 83 | 67.3 ± 27 | 63.7 ± 20 |
|  | 84 | 106.0 ± 38 | 66.8 ± 23 |

**Table 2.**
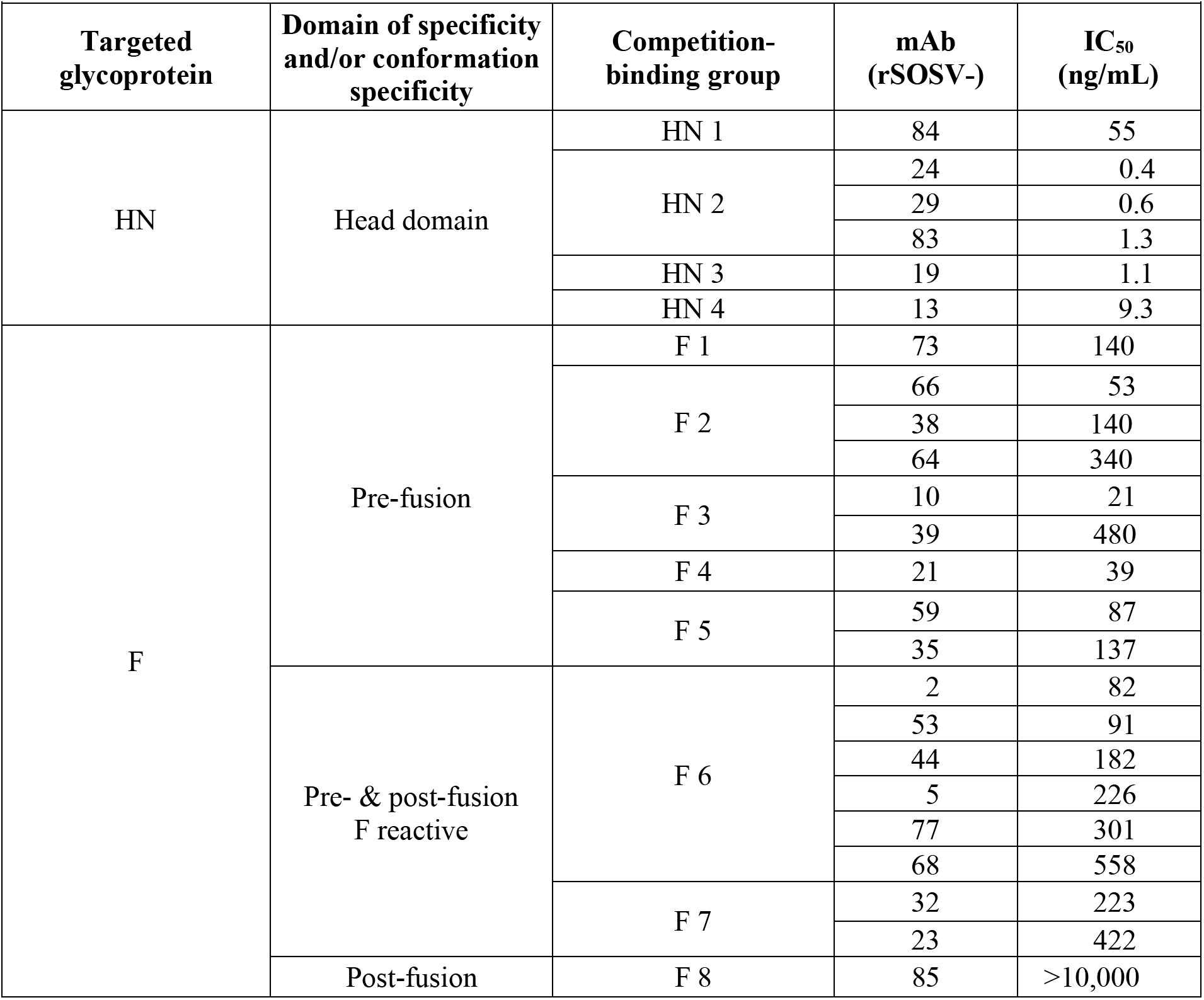
Half-maximal inhibitory concentration (IC_50_) values for SOSV HN- or F-reactive mAbs in neutralization assay using authentic SOSV.

### Domain specificity of HN-reactive mAbs

The specificity of the HN-reactive mAbs was refined further by testing for binding of the rmAbs to the recombinant HN_head_ protein or the tetrameric/dimeric HN_ecto_ protein in ELISA. All six of the HN-reactive mAbs (SOSV-13, −19, −24, −29, −83, and −84) bound to both HN antigens (**Figure 3**), indicating that the binding sites for all mAbs were likely located in the globular head domain of the HN protein. The half-maximal effective concentration (EC_50_) values for binding of the rmAbs to soluble HN proteins ranged from 22 to 92 ng/mL (**Table 1**). For comparative purposes, two of the mAbs (SOSV-84 and SOSV-24) were purified as IgG from hybridoma cell line supernatants and tested in ELISA and had similar EC_50_ values for binding compared to their respective recombinant versions (not shown). The HN-reactive rmAbs also were tested in a competition-binding ELISA (**Figure 4**). The data revealed that there are four HN competition-binding groups represented in the panel with members as follows, <u>Group 1:</u> SOSV-84, <u>Group 2:</u> SOSV-83, SOSV-29, and SOSV-24, <u>Group 3:</u> SOSV-19, and <u>Group 4:</u> SOSV-13. Four competition-binding groups were identified consistently, whether testing for competition-binding to HN_head_ or HN_ecto_ proteins (**Figure 4**).

**Figure 3.**
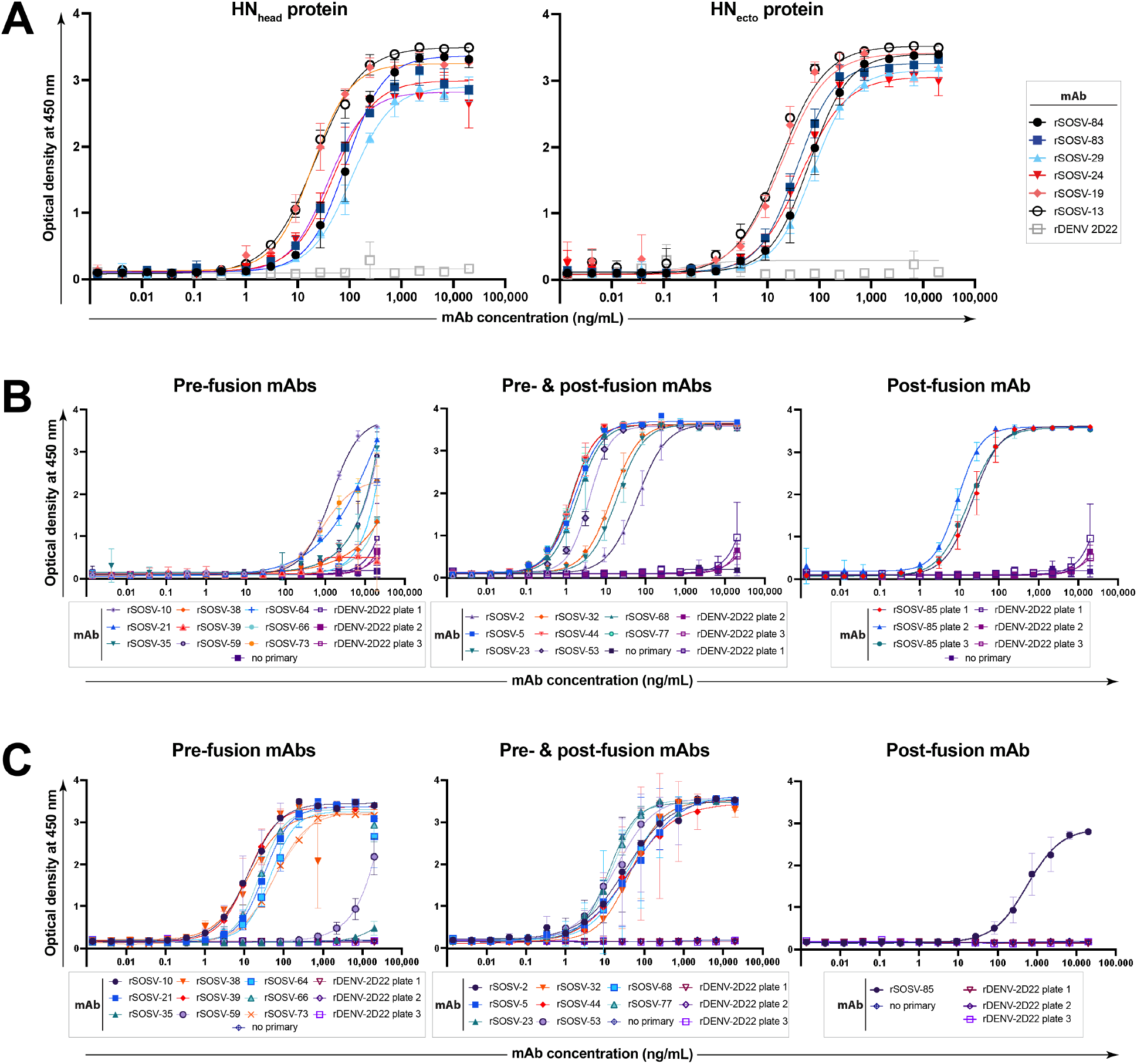
Representative curves in ELISA for binding of rSOSV mAbs to soluble glycoprotein antigens. Shown here is a single, representative set of data of the three biological replicates that were performed. Curves show the average of three technical replicates plotted with standard deviation. The anti-F rmAbs are divided into 3 subsets: pre-fusion, pre- & post-fusion, or post-fusion, although all the rmAbs were tested simultaneously. **(A)** Anti-HN rSOSV mAbs against HN_ecto_ and HN_head_ proteins. **(B)** Anti-F rSOSV mAbs against post-fusion SOSV F. **(C)** Anti-F rSOSV mAbs against SOSV pre-fusion F construct.

**Figure 4.**
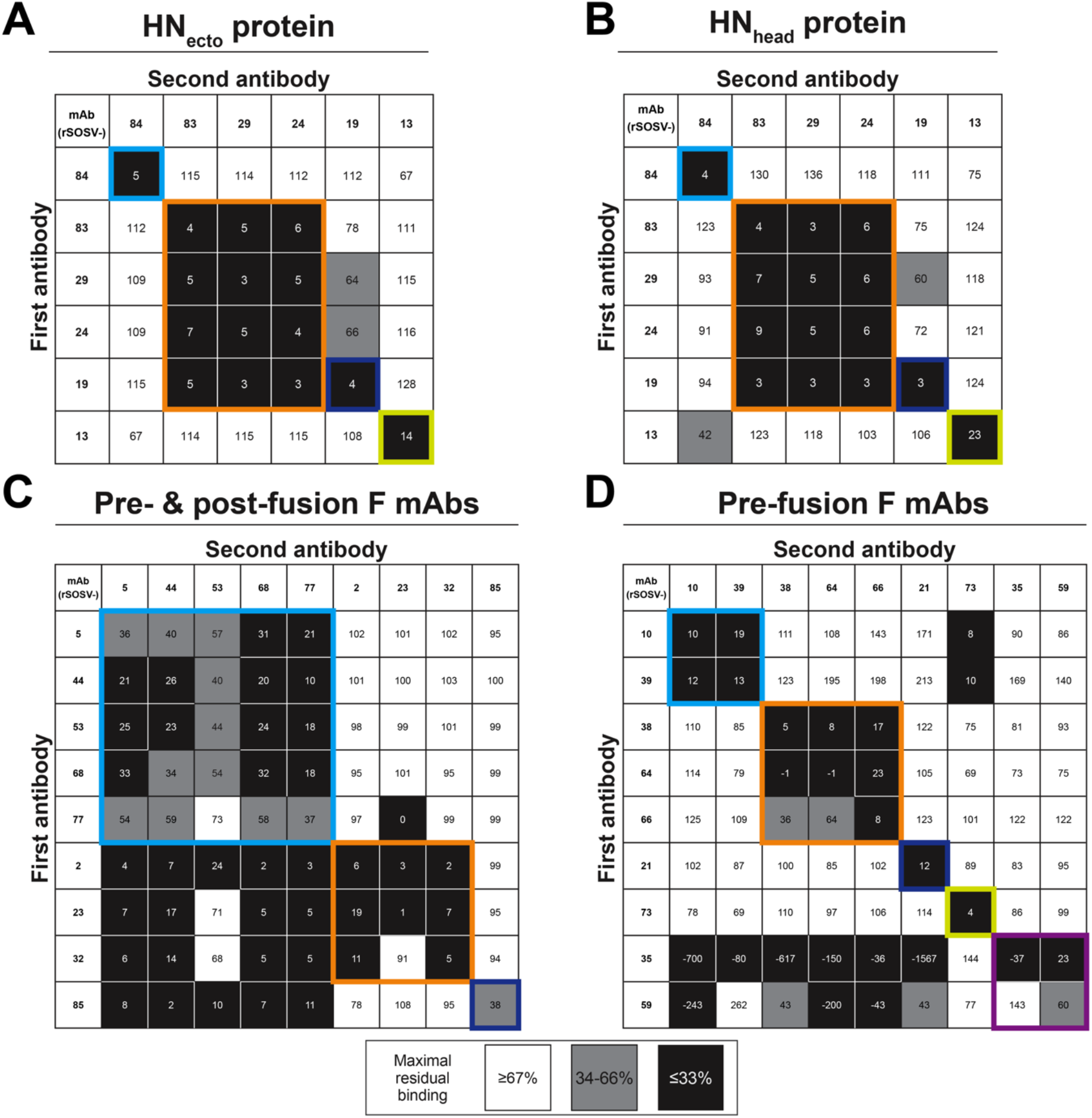
Competition-binding assay for anti-HN mAbs. Unlabelled primary (first) mAb was bound to antigen-coated plates in saturating conditions (10 μg/mL) with secondary antibodies added at a final concentration of 100 ng/mL for HN-mAbs and 500 ng/mL for F-mAbs according to the grid layout shown. Data were converted to percent binding relative to the maximal un-competed binding of the second antibody (lacking a primary mAb). Assays were repeated in triplicate with quadruplicate technical replicates. A representative assay for each antigen/mAb set tested is shown. **(A)** Binding data of mAbs against HN_ecto_ as antigen. **(B)** Binding data of mAbs against HN_head_ as antigen. **(C)** Binding data for pre- & post-fusion anti-F mAbs (rSOSV-2, 5, 23, 32, 44, 53, 68, & 77) and the post-fusion mAb (rSOSV-85) against pre-fusion F protein. **(D)** Binding data for pre-fusion F specific mAbs (rSOSV-10, 21, 35, 38, 39, 59, 64, 66, & 73) against pre-fusion F antigen.

**Figure 5.**
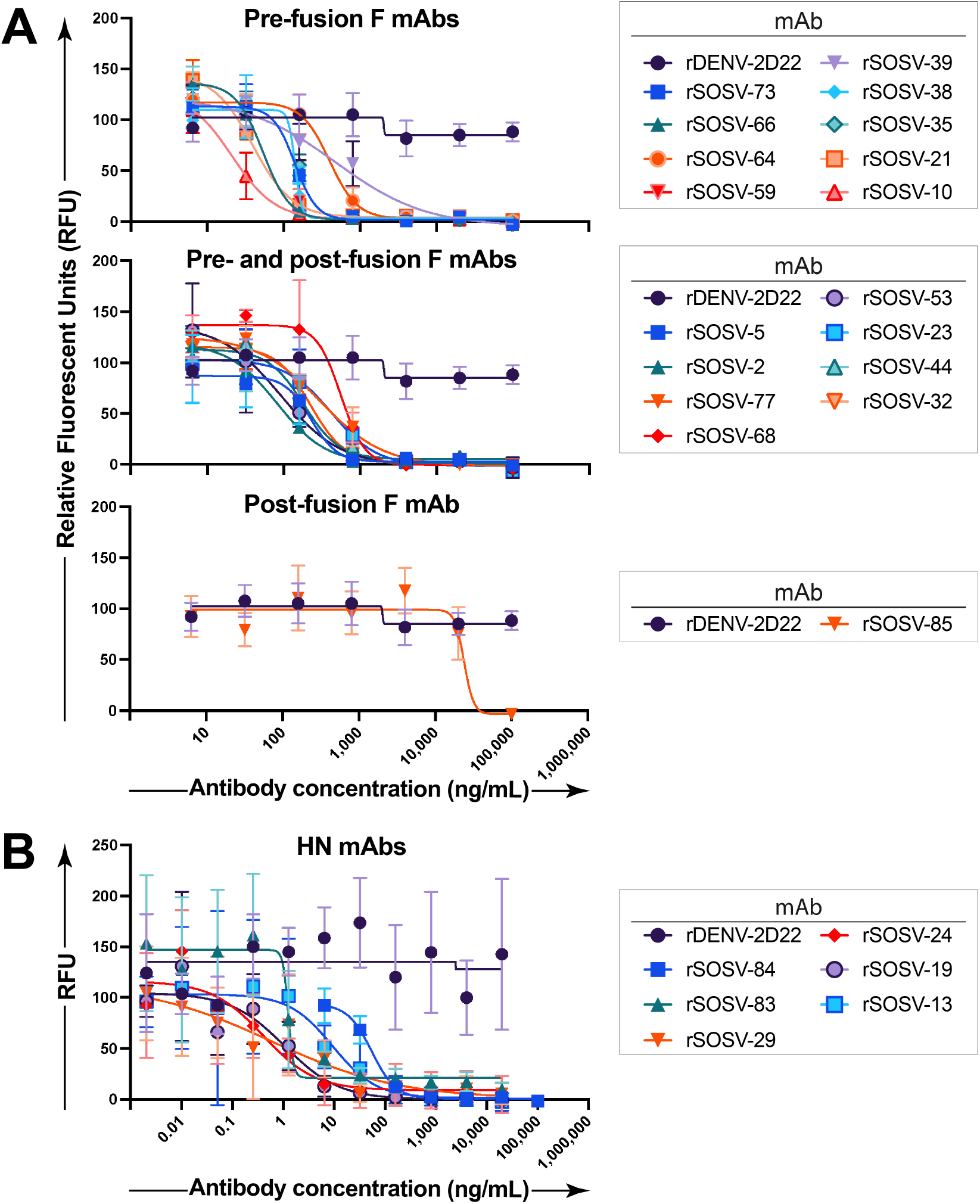
Neutralization assay of SOSV mAbs against live virus. SOSV mAbs were tested for inhibition of authentic rSOSV-ZsG in quadruplicate on Vero-E6 cell culture monolayers. **(A)** Neutralization data for anti-F mAbs. Data are grouped according to the pattern of antigen-reactivity: pre-fusion F, pre- and post-fusion F, or post-fusion F protein. **(B)** Neutralization data for the HN-specific mAbs.

### Conformational specificity of anti-F mAbs

Since paramyxovirus fusion proteins are well known to only be metastable in the pre-fusion state (11, 23), we used soluble post- and pre-fusion-stabilized proteins to test for conformational specificity of the panel of anti-F mAbs. Only one mAb (rSOSV-85) was specific to the post-fusion conformation while seven mAbs (SOSV-10, −21, −38, −39, −64, −66, and −73) were specific to the pre-fusion conformation (Figure 3). Eight mAbs (SOSV-2, −5, −23, −32, −44, −53, −68, −77) recognized both the post- and pre-fusion proteins. Two mAbs, rSOSV-35 and rSOSV-59, did not bind detectably to either pre- or post-fusion soluble F protein in ELISA (**Figure 3**) although they did bind to cell-surface displayed, WT protein. We theorize that the WT F protein is predominantly in the pre-fusion state due to the absence of co-expression of the HN protein (11, 23–25) and the fact that so few post-fusion specific mAbs were isolated in our cell-surface display screening. Thus, it is likely that rSOSV-35 and rSOSV-59 are specific for pre-fusion F, and so we included them in this group. The half-maximal effective concentration (EC_50_) values for binding of the rmAbs are summarized in **Table 1** and range from 1.5 ng/mL to over 4,000 ng/mL against post-fusion F and 15.7 ng/mL to over 1,000 ng/mL for pre-fusion F. As with the anti-HN mAbs, we further categorized the anti-F mAbs by competition-binding ELISA (**Figure 4**) on their respective SOSV F conformational specificity. The data revealed that there are approximately eight distinct competition-binding groups: <u>Group 1)</u> SOSV-73; <u>Group 2)</u> SOSV-66, −64, and −38; <u>Group 3)</u> SOSV-39 and SOSV-10; <u>Group 4)</u> SOSV-21; <u>Group 5)</u> SOSV-59 and SOSV-35; <u>Group 6)</u> SOSV-77, −68, −53, −44, −5; <u>Group 7)</u> SOSV-32, −23, −2; and <u>Group 8</u>) SOSV-85.

### Neutralizing activity of human mAbs against authentic SOSV

We next tested each of the 24 human mAbs for neutralization against recombinant SOSV expressing ZsGreen (26) by incubating dilutions of mAb with SOSV and adding to Vero cell culture monolayers. Quantification of the ZsGreen reporter was used to quantitate SOSV infection in the monolayer and calculate the IC_50_ value for each mAb. Each of the mAbs inhibited virus infection, except for the post-fusion specific mAb. As a class, the HN-specific mAbs neutralized more potently than the anti-F mAbs. Of the anti-F panel, the pre-fusion-specific mAbs neutralized more potently. Even though this individual experienced a primary infection with a novel paramyxovirus, the infection induced some antibodies with ultra-potent (<1 ng/mL) neutralizing activity, which are some of the most potently neutralizing human antibodies to a virus ever reported.

## Discussion

These studies of human immunity to SOSV are of interest because they elucidate the molecular basis for response to an emerging virus in the *Pararubulavirus* genus, which is understudied. The global virome, especially the virome of bats in the wild, contains many viruses that have epidemic potential (27), but paramyxoviruses are perhaps the most common threats (28). Little is known of the determinants of immunity for most of these viruses. There is a licensed vaccine for mumps virus, which is a prototype virus in the *Orthorubulavirus* genus, but the mechanisms by which this vaccine protects and the correlates of protection against mumps virus infection and disease are not defined. Here we surveyed the antigenic landscape on the two major surface proteins of SOSV to understand the principal determinants of immunity for a virus in this genus. We isolated naturally occurring human mAbs that recognized HN or F proteins since other paramyxoviruses’ attachment and fusion proteins usually contain sites of vulnerability for neutralization. Before this work, it was not clear if SOSV HN and F proteins induce neutralizing antibodies, and if so, what is the more immunodominant protein, and what are the regions on these proteins inducing neutralizing antibodies.

Of the six anti-HN mAbs isolated, each recognizes sites on the globular head domain of HN. Even though all the HN-reactive mAbs target the head domain, the competition-binding patterns reveal that there are at least four distinct major antigenic sites recognized by these mAbs. Most of the antibodies we isolated from this individual neutralized authentic SOSV, and some of them did so with very potent activity (exhibiting very low IC_50_ values in *in vitro* neutralization assays). Since the HN protein is the principal attachment factor for paramyxoviruses, anti-HN SOSV mAbs might be expected to possess more potent neutralizing activity than F-protein-specific mAbs, and indeed we found this to be the case. The high neutralizing potency of the Group 2 anti-HN mAbs suggests that these antibodies may block HN protein binding to cell surface receptors. It is possible that the prolonged and marked viremia induced ultra-potent mAbs may have contributed to the survival of the in this wildlife researcher but might not be elicited as commonly in humans during mild infection. An unanswered question is the seroprevalence of SOSV in people living in South Sudan or Uganda where the Egyptian rousette bats are present, or whether if SOSV infection in people living in areas endemic with pararubulaviruses would have the same susceptibility to severe disease as this U.S.-origin wildlife researcher. The receptor for SOSV is not known, but future studies of these mAbs may help in efforts to discover the receptor for SOSV and possibly related pararubulaviruses. Blocking attachment is often one of the most efficient and complete mechanisms of antibody-mediated neutralization. However, this is not always the case since antibodies to pneumovirus fusion proteins often are more potent as a class than antibodies to the attachment protein for those viruses (for example, antibodies to respiratory syncytial virus or human metapneumovirus F and G surface glycoproteins) (29).

The F protein of paramyxoviruses can exist in two very distinct conformations, pre-fusion and post-fusion. Most post-fusion-specific F antibodies are expected to be non-neutralizing or poorly neutralizing since virions particles possess the pre-fusion form of F, and the post-fusion state occurs following viral fusion with host cell membranes. The pre-fusion conformation of the F protein, however, is a potential neutralizing target since antibodies that block the triggering of the pre-fusion F protein may prevent virus-cell fusion. We tested our F-specific mAbs for their ability to recognize pre- or post-fusion F proteins and found that we could classify our panel into three broad groups: mAbs that recognized 1) pre-fusion, 2) post-fusion, or 3) both conformations. As expected, the pre-fusion-specific mAbs as a group tended to have higher neutralization potency (*i.e*., lower IC_50_ values) compared to the post-fusion-specific mAb and many of the pre- and post-fusion specific mAbs. Of the panel of 24 mAbs, only the post-fusion specific mAb failed to neutralize virus. The SOSV-survivor’s potent antibody response at 5 years post-after infection (presumably without re-exposure to virus while living in the U.S.) to both F and HN glycoproteins is remarkable and suggests the possibility for inducing durable vaccine-induced durable functional immunity in humans should the need ever arise.

This panel of mAbs may be useful in several applications. An ultra-potent HN mAb, such as rSOSV-24, is potentially a therapeutic candidate given its extraordinarily low IC_50_ value for neutralization of 0.4 ng/mL. As there are currently no available SOSV-specific reagents, the mAbs discovered in this work also can serve as reagents for the continued study of SOSV pathogenesis and immunity. rSOSV-85 as a post-fusion-specific mAb can be used in various applications, such as to study the fusion-triggering process during the SOSV life cycle, help test the stability of potential pre-fusion F protein vaccine candidates, or aid in pre-fusion F protein purification processes by sequestering post-fusion F protein during chromatographic purification protocols. Potently neutralizing pre-fusion-specific F mAbs like rSOSV-10 can serve as positive controls in neutralization assays for testing anti-viral compounds or vaccines. Finally, knowledge of the competition-binding groups of the HN and F proteins may help in further understanding protein domains governing the paramyxovirus fusion process or in receptor discovery studies for SOSV. As SOSV and other pararubulaviruses lack the ability to bind to sialic acid but can infect human cells (15, 16), discovering the receptor for this genus of paramyxoviruses could greatly advance efforts for epidemic preparedness against this group of viruses. Also, it is quite likely that the receptor is conserved between bats, humans, and pigs, as evidenced by Teviot, Tioman, and Menangle viruses (4, 6, 15, 30, 31)—indicating potential threat to and from domestic livestock as well.

In summary, the human mAbs isolated in this study are the first SOSV-specific mAbs generated and can be used for further studies of SOSV and related viruses and as candidate therapeutic antibodies for clinical development.

## Methods

### Immune cells

In 2012, a 25-year-old, otherwise healthy individual was infected with SOSV during occupational exposure while handling wild bats as part of wildlife surveillance activities in South Sudan and Uganda. Clinical and virologic features of infection for this only known human survivor of SOSV infection have been described previously (7). RNA extraction from the individual’s whole blood followed by RT-PCR amplification and DNA sequence analysis revealed the presence of RNA genome for a novel paramyxovirus that was named *Sosuga pararubulavirus* (or Sosuga virus [SOSV]) after the countries visited by the wildlife researcher (<u>So</u>uth <u>S</u>udan and <u>Uga</u>nda) (7). A leukapheresis pack was obtained from the individual after written informed consent in 2017, approximately five years after the infection. Peripheral blood mononuclear cells (PBMCs) were isolated from the leukapheresis product, cryopreserved at a density of 10 or 25 million cells/mL, and stored in the vapor phase of liquid nitrogen until use. The studies were approved by the Vanderbilt University Medical Center Institutional Review Board.

### EBV transformation of cell lines from human blood

Vials of cryopreserved PBMCs were thawed at 37°C and washed in ClonaCell™-HY Medium A (STEMCELL Technologies, cat. #03801). Epstein-Barr virus (EBV) was obtained by collecting the supernatant of the marmoset lymphoblastoid cell line (LCL) B95-8 (32) (formerly available from the American Type Culture Collection as ATCC^®^ CRL-1612). B cells were transformed with EBV by combining washed PBMCs with prepared stocks of filtered B95-8 cell supernatant, using 4.5 mL to transform 8 to 10 million PBMCs in B cell growth medium (Medium A containing CpG [Invitrogen, oligo ZOEZOEZZZZZOEEZOEZZZT] at the 10 μmole scale [desalted], cyclosporin A [Sigma-Aldrich, cat. #C1832], and Chk2 inhibitor II [Sigma-Aldrich, cat. # C3742]). Cells were plated at 50 μL/well in one 384-well plate for each suspension of 8 to 10 million PBMCs. Cells were incubated at 37°C in 7% CO_2_ for 6 to 12 days until LCLs were clearly visible and forming colonies. The plates of transformed B cells were expanded to four 96-well plates in B cell expansion medium (Medium A, CpG, Chk2i II, and 10 million irradiated human PBMCs per plate from an unrelated healthy donor [Nashville Red Cross]). The plates were incubated at 37°C in 7% CO_2_ for 4 to 7 days before screening antibodies in LCL supernatants for binding to SOSV antigen expressed in cells using a high-throughput flow cytometry assay.

### Production of human hybridoma cell lines from transformed B cells

Once positive SOSV-reactive wells of LCLs were identified, the B cells from these wells were transferred to microcentrifuge tubes and washed three times with BTX medium (300 mM sorbitol [Fisher Scientific, cat. # BP439], 0.1 mM calcium acetate [Fisher Scientific, cat. # AC21105-2500], 0.5 mM magnesium acetate [Fisher Scientific, cat. #AC42387-0050], and 1.0 mg/mL bovine serum albumin [Sigma-Aldrich, cat. # A3294]). The B cell pellets were resuspended in BTX medium, combined with the HMMA2.5 human-mouse myeloma fusion partner cell line and electroporated in a 0.2 μm cuvette (BTX, cat. # 45-0125). After fusion, cells were left in cuvettes in 7% CO_2_ at 37°C for at least 30 min before transferring to hypoxanthine-aminopterin-thymidine (HAT) selection medium (20% ClonaCell™-HY Medium E [STEMCELL, cat. #03805], 80% Medium A, HAT media supplement [final concentrations: 100 μM hypoxanthine, 0.4 μM aminopterin, 16 μM thymidine (Sigma-Aldrich cat. # H0262-10VL)], and 150 μL of 1 mg/mL ouabain octahydrate [Sigma-Aldrich, cat. #O3125; final concentration 0.33 μg/mL,]). Fused cells were plated by limiting dilution in 384-well plates with 50 μL/well volumes. The plates were incubated for 2 to 3 weeks, feeding with 25 μL/well of Medium E after 1 week, before screening for binding to recombinantly expressed viral antigens by high-throughput flow cytometry to identify wells with hybridomas secreting SOSV-reactive antibodies. SOSV-reactive hybridomas were expanded to 48-well plates with 500 μL/well Medium E and screened again.

### Isolation of human mAbs secreted from hybridoma cell lines

The hybridoma cell lines secreting SOSV-reactive antibodies were cloned using single-cell sorting on a BD FACSAria III cytometer (BD Biosciences) or SH800 Cell Sorter (Sony Biotechnology) into 384-well plates containing Medium E and incubated in 7% CO_2_ at 37°C for 1 to 2 weeks for the cells to expand in number. Supernatants from 384-well plates (one plate for each hybridoma line) were screened by high-throughput flow cytometry using cell-surface expressed SOSV antigens to identify hybridoma cell clones secreting SOSV-specific antibodies. The cloned hybridoma cell lines with antigen-reactive supernatants were scaled up gradually in 48-well, 12-well, T-25, and T-75 plates or flasks with screening for antibody binding to cell-surface displayed antigens by high-throughput flow cytometry at each expansion step. The cells from T-75 flasks were used to make frozen stocks of the mAb-secreting cloned hybridoma cell lines by freezing cells in freezing medium (50% cell culture medium, 40% fresh Medium E, and 10% dimethyl sulfoxide).

### Sequence analysis of antibody variable genes

Cell pellets of the cloned hybridoma cell line cultures were processed for RNA extraction and amplification of antibody variable genes by 5’RACE or 3’RACE procedures, and DNA sequence analysis of cDNA using a Sequel instrument (Pacific Biosciences) as previously described (33).

### Purification of mAb proteins

MAb IgG proteins in supernatants of cloned hybridoma cell lines were prepared by washing cells from T-75 flasks in serum-free Hybridoma-SFM Medium (Thermo Fisher Scientific, cat. # 12045076) and seeding 3 to 6 wells of a 6-well G-Rex plate (Wilson Wolf, cat. # 80240M) with the mAb-secreting lines in Hybridoma-SFM medium. The G-Rex plates were incubated in 7% CO_2_ at 37°C, with the supernatant typically being harvested and cells split every one to two weeks. The G-Rex wells were reseeded up to a maximum of 3 times. MAb supernatants were collected and clarified through a 0.2 μm filter, and then mAbs were isolated by fast protein liquid chromatography (FPLC) on an ÄKTA pure system (Cytiva) using HiTrap Protein G High Performance (Cytiva, cat. # 17-0404-01) or HiTrap MabSelect SuRe (Cytiva, cat. # 11-0034-95) columns. Soluble forms of the SOSV antigens with were purified by using the proteins’ 6×-His tag or StrepII tags and Expi293F cells and Expifectamine 293 expression system (Thermo Fisher Scientific) as discussed below at day 5 to 7 post-transfection. Transfected cell supernatant was collected, and the 6×-His- or StrepII-tagged antigens purified using the ÄKTA pure system with HisTrap™ Excel (Cytiva, cat. # 17-3712-05), StrepTrapHP™ (Cytiva, cat. # 29-0486-53), or StrepTrapXT™ (Cytiva cat. # 29401322) columns as appropriate. Eluates were run on an SDS-PAGE gel to identify samples with the target proteins. These samples were collected and concentrated using an Amicon Ultra-15 Centrifugal Filter with Ultracel-10 Membrane (Millipore Sigma, cat. # UFC901024) and buffer exchanged to DBPS with Zeba™ Spin Desalting Columns (7K MWCO, 10 mL, Thermo Fisher Scientific, cat #89894).

### High-throughput flow cytometric detection of binding to cell-associated viral antigens

Expi293F cells were transfected with DNA plasmids encoding either wild-type full-length SOSV F or HN constructs. Cells were seeded in flat-bottomed flasks at 2.5 × 10^6^ cells/mL, using the volume of the culture size used to scale the transfection mix. The transfection mix was prepared by combining cold Opti-MEM™ I Reduced-Serum Medium (Thermo Fisher Scientific; 0.1 mL/mL of cells), DNA (1 μg/mL of cells), and 2.7 μL/mL of cells Expifectamine 293 reagent (in Thermo Fisher Scientific kit, cat. # A14524) mixing 3 to 5 times by pipetting and incubating at RT for 20 to 30 min. Flasks of cells first were swirled before adding the transfection mix to ensure even spreading of the transfection mix. Cells were incubated at 37°C in 7% CO_2_ with shaking at 125 RPM for 24 to 48 hr. The day after transfection, ExpiFectamine™ 293 Transfection Enhancer 1 and Enhancer 2 were added at 0.5% or 5% scale of transfection, respectively. Transfected cells were plated at 50,000 to 70,000 cells/well in 96-well V-bottom plates, washed with flow cytometry buffer (DPBS without calcium and magnesium, 2% low-IgG FBS, 2 mM EDTA), and stained with antibodies in the supernatants of transformed B cells or hybridoma cells, or purified mAb, at 30 to 50 μL/well at 4°C for 30 min as the primary stain. The primary stain was washed off with flow cytometry buffer, and the cells were stained with 50 μL/well of a 1:1,000 dilution of goat anti-human IgG-PE (Southern Biotech, cat. # 2040-09) secondary antibodies for 30 min at 4°C. The secondary antibodies were removed by washing with DPBS and cells were fixed with 50 to 100 μL of 4% paraformaldehyde (PFA) in DPBS for 10 min at RT. For screening for binding of antibodies in hybridoma supernatants or suspensions of purified mAbs, cells were stained with LIVE/DEAD™ Fixable Violet Dead Cell Stain (Invitrogen, cat. # L34963) for 30 min at 4°C prior to fixation. The fixative was washed off with flow cytometry buffer and cells resuspended in 25 μL of that buffer and analyzed on an iQue Screener PLUS cytometer (Sartorius).

### Soluble SOSV F and HN protein constructs

Coding sequences for the wild-type SOSV F or HN proteins were obtained from the 2012 human isolate sequences (GenBank NC_025343.1), sequence-optimized for human expression, and cDNA was synthesized by Twist Bioscience and inserted into pTwist-CMV expression vectors for use in cell-surface expression assays. Additionally, constructs of the HN and F wild-type coding sequences were generated with the same sequences as above but with the addition of cDNA encoding a DYKDDDDK (FLAG^®^) tag on the cytoplasmic domain of the proteins (carboxy terminus for the F protein and amino terminus for the HN protein) and synthesized by Twist Bioscience. Plasmids containing cDNAs encoding soluble forms of the HN protein were synthesized by Twist Bioscience with a CD5 signal peptide sequence for secretion and a thrombin-cleavable 6× His tag for purification. The construct for the ectodomain of the HN protein (designated HN_ecto_) contains amino acid residues 75 to 582, while the HN head domain construct (designated HN_head_) contains residues 125 to 582, as previously described (16). Soluble forms of the pre-fusion F protein were designed that included residues 15 to 476 of the SOSV F sequence with the following modifications: amino acid changes I206C, A223C, K101-F103GGG were introduced, a GCNt domain was added to the C-terminus, and a mouse IL-2 signal peptide was placed into the pVRC8400 vector at the N-terminus of the protein-coding sequence. The I206C and A223C mutations create a disulfide bond, and K101-F103GGG edits the furin cleavage site.

### Microscopy of SOSV F and HN expression in cultured cells

HEK293T/17 (ATCC, cat #CRL-11268) cells were seeded at 20,000 live cells/well into a clear-bottomed, black 96-well plate (Greiner Bio-One, cat. #655090) in DMEM + 10% FBS + 1% PSG. While in suspension, cells were transfected with 10 μL of Lipofectamine 3000 (Thermo Fisher Scientific, cat. #L3000015) transfection mix containing plasmids encoding SOSV F-WT, SOSV HN-WT, SOSV F-WT & SOSV HN-WT, SOSV F-FLAG & SOSV HN-FLAG, VSV-G, or no DNA (mock), with 12 replicate wells for each condition. Transfection mixes were prepared following the manufacturer’s protocol using ~0.15 μL of Lipofectamine 3000 reagent and ~100 ng total DNA per well. The plate was incubated at 37°C in 5% CO_2_ for 45 hr, after which the medium was removed, and the cells were fixed in 100 μL of 4% PFA for 1 hr. Cells were washed several times with DPBS before blocking and permeabilizing with 150 μL/well of permeabilization buffer (5% milk, 0.1% saponin, in 1× PBST) or 60 min at room temperature (RT). A polyclonal mix of anti-SOSV mAbs was made by mixing 3 HN mAbs and 3 F mAbs; the anti-SOSV mix was diluted to 1:250 in permeabilization buffer while monoclonal anti-FLAG M2 (Sigma-Aldrich, cat. # F3165) was diluted 1:500 in permeabilization buffer. Cells were stained with 50 μL of anti-SOSV or anti-FLAG primary (half the plate) and incubated ~1 hr at RT, after which the primary stain was removed, and the plate washed with DPBS. The cells were then stained with 50 μL of the secondary mix (goat anti-human IgG-AF488 (SouthernBiotech, cat. #2040-30) and goat anti-mouse Alexa Fluor 488 IgG (H + L) (Invitrogen, cat. #A11001) both diluted to 1:1,000) in permeabilization_buffer and incubated 1 hr at RT, protected from light. Cells were then washed with DPBS before being stained with 50 μL/well of DAPI (4’,6-diamidino-2-phenylindole, dihydrochloride) (Invitrogen, cat. #D1306) diluted to 5 μM in DPBS for 15 min. Cells were then washed several times and then kept in 250 μL/well of DPBS for imaging. Imaging was done on an EVOS M5000 instrument (Invitrogen, cat. #AMF5000) with a 10× objective with four fields of view imaged for 3 replicate wells of each transfection condition. Area of stained cells/syncytia was measured using Fiji (34), and the data were analyzed in Prism (GraphPad Software, version 9.3.1 for Mac OS X).

### Enzyme-linked immunosorbent assay (ELISA) to detect antibody binding to viral proteins

ELISAs were performed by coating 384-well plates with either soluble viral glycoprotein (HN_ecto_, HN_head_, pre-fusion F with 6× His and StrepII tags [preF-tHS], or post-fusion F with 6× His and StrepII tags [postF-tHS] at 2 μg/mL in 20 to 25 μL of DPBS. Plates were coated with antigen overnight at 4°C, washed three times with phosphate buffered saline with Tween 20 (PBS-T; Cell Signaling, cat. #9809S, 20× stock solution used to make 0.05% Tween 20 when diluted to 1×) using an EL406 plate washer dispenser instrument (BioTek), then blocked for 1 to 3 hr at RT with 50 μL/well of blocking buffer: 2% Blotting Grade Blocker (Bio-Rad cat. # 1706404) and 2% heat-inactivated goat serum (Gibco, cat. #16210-072) in PBS-T. SOSV HN mAbs were diluted in blocking buffer starting at 20 μg/mL in a 3-fold serial dilution series. After removing the blocking buffer, primary antibodies were added at 20 μL/well to the plates and incubated at RT for 1 hr. Plates were washed three times with PBS-T prior to addition of the secondary antibodies. The secondary antibody solution was prepared by diluting goat anti-human IgG HRP-conjugated antibodies (SouthernBiotech, cat. # 2040-05) at 1:2,000 in blocking buffer and adding 20 to 25 μL/well, and then incubated at RT for 1 hr. Secondary antibodies were removed, and plates washed three times with PBS-T. A volume of 25 μL/well of 1-step Ultra TMB-ELISA Substrate Solution (Thermo Fisher Scientific, cat. # PI34029) was added to the plates and incubated for 5 to 10 min at RT, before being quenched with 25 μL of 1N hydrochloric acid (Fisher Scientific, cat. # SA48-1). Plates were analyzed on a BioTek plate reader at 450 nm wavelength. Data were analyzed in Prism (GraphPad Software, version 9.3.1 for Mac OS) using a sigmoidal, four-parameter logistic, nonlinear regression model to generate the graphs and EC_50_ values for the mAbs.

### Biotinylation of SOSV-specific antibodies

SOSV F- or HN-reactive mAbs and a similarly prepared human mAb (rDENV-2D22) specific for an unrelated virus antigen (dengue virus envelope protein) were biotinylated. Purified IgG mAb proteins were diluted to a concentration of 1 mg/mL in DPBS, and an aliquot of 200 μL volume (containing 200 ng of antibody) was used for biotinylation. A 2 mg vial of EZ-Link™ NHS-PEG4-Biotin, No-Weigh™ Format biotin (Thermo Fisher Scientific, cat. #A39259) was reconstituted with 170 μL of DPBS or dimethyl sulfoxide and 1.33 μL of the biotin solution and added to 200 ng of each of the purified antibodies. Antibody-biotin solutions were mixed and incubated at RT for 50 min. Excess biotin was removed using Zeba™ Spin Desalting Plates, 7K MWCO (Thermo Fisher Scientific, cat. #89807). The plate columns were equilibrated with DPBS following the manufacturer’s protocol. The antibody-biotin mixtures were loaded onto two columns for each mix, with ~100 μL loaded onto each column. The resulting duplicate eluates were combined.

### Competition-binding studies using ELISA

Competition ELISAs for the anti-SOSV mAbs were performed by coating 384-well plates overnight at 4°C with 20 μL of 2 μg/mL concentration solutions of antigen in DPBS: HN_ecto_ or HN_head_ protein for anti-HN mAbs and pre-fusion or post-fusion F protein for anti-F mAbs. Plates were washed three times with PBS-T using an EL406 plate washer (BioTek), then blocked for 1 to 3 hr at RT with 50 μL/well of blocking buffer (HN: 5% Blotting Grade Blocker, Bio-Rad cat. # 1706404 in PBS-T or F: 2% Blotting Grade Blocker and 2% goat serum, Gibco, cat. #16210-072 in PBS-T). Blocking buffer was removed by washing plates three times with PBS-T on an EL406 plate washer. The SOSV mAbs or control mAb DENV-2D22 were diluted to a concentration of 10 μg/mL in respective blocking buffers, and 20 μL of each mAb was plated into wells of a 384-well plate to give quadruplicate readings for each mAb combination. To determine the maximal binding of each mAb in the absence of competition, 20 μL of plain blocking buffer (without a primary antibody) was placed into enough wells of a 384-well plate to give quadruplicate readings for each mAb combination. Plates were incubated at RT for 1 hr. The biotinylated HN-mAbs were diluted to 500 ng/mL in blocking buffer while the F-mAbs were diluted to 2,500 ng/mL in blocking buffer, and 5 μL of biotinylated mAb was added to the 20 μL of unlabelled mAb or blocking buffer control, so that the final concentration of biotinylated antibody was 100 ng/mL for anti-HN mAbs and 500 ng/mL for anti-F mAbs. Plates were incubated at RT for 1 hr, and then were washed three times with PBS-T using an EL406 plate washer. A volume of 25 μL of a 1:1,000 dilution of Mouse Anti-Biotin-AP (Southern Biotech, cat. #6404-04) in blocking buffer was added to each of the wells of the HN plates and incubated for 1 hr at RT. For the F-coated plates, 25 μL of a 1:2,000 dilution avidin-peroxidase (Sigma-Aldrich, cat. # A7419-2ML) in blocking buffer was added to each of the wells of the plates and incubated for 1 hr at RT. The AP-labelled antibody or avidin-peroxidase was removed with three washes of PBS-T. For HN plates, 25 μL of phosphatase substrate (Sigma-Aldrich, cat. # S0942) was diluted to 1 mg/mL in AP-substrate buffer (pH 9.6, 1M Tris Base [Tris (Hydroxymethyl) Aminomethane]; Research Products International, cat. # T60040), 0.3 mM MgCl_2_ [Sigma-Aldrich, cat. # M1028]) and added to each well. Plates were developed in the dark at RT for 1 hr before being read on a BioTek plate reader at 405 nm wavelength. For the F plates, 25 μL/well of 1-step Ultra TMB-ELISA Substrate Solution (Thermo Fisher Scientific, cat. # PI34029) was added to the plates and incubated for 5 to 10 min at RT, before being quenched with 25 μL of 1N hydrochloric acid (Fisher Scientific, cat. # SA48-1) and read on BioTek plate reader at 450 nm wavelength. Since some anti-F mAbs (rSOSV-10, 21, 35, 38, 39, 59, 64, 66, & 73) had shown poor binding to post-fusion F, these mAbs were not tested for competition-binding on the post-fusion F protein. However, anti-F mAbs rSOSV-2, 5, 23, 32, 44, 53, 68, 77, and 85 were tested on both pre-fusion and post-fusion F. The values obtained from quadruplicate wells were averaged, and values from the wells with the negative control mAb rDENV-2D22 were considered the nonspecific binding signal and subtracted. The averaged absorbance data then was converted to percentage relative to the maximal (without unlabelled primary mAb) data. Competing mAbs were defined as having a residual binding level equal to or below 33% of the maximal binding level, intermediate competition was defined as having 34 to 66% of the maximal binding, and non-competing mAbs were defined as having equal to or greater than 67% of maximal binding.

### Neutralization of SOSV by human mAbs

The anti-SOSV mAbs were tested for neutralization activity with authentic virus under biosafety level 3 (BSL-3) conditions. To assist in viral quantification, we used a previously described recombinant SOSV encoding ZsGreen (ZsG) protein (26). Neutralization activities of mAbs were measured using a standard protocol in Vero-E6 cell monolayer cultures. Briefly, serial five-fold dilutions of mAbs (150 μL) made in Dulbecco’s Modified Eagle Medium (DMEM) were mixed with an equal volume of a suspension of SOSV-ZsG containing 100 median tissue culture infectious dose (TCID_50_). After incubation at 37°C for 1 hr, 50 μL of virus-antibody mixture was inoculated onto each well containing a Vero-E6 cell monolayer culture in 96-well plates (Cell Carrier Ultra plates, Perkin Elmer) that had been seeded the day before at 15,000 cells per well. The culture was incubated at 37°C for 72 hr, and after that fluorescence intensities were determined using a multi-well plate reader (Synergy; BioTek). Fluorescence readings were taken from quadruplicate wells at each mAb concentration. Background fluorescence signals (obtained from wells lacking virus) were deducted from the virus and treatment readings, and data are presented as the percent of the no-antibody and virus-only control. Prism software (GraphPad) was used to generate concentration–response plots. A four-parameter equation was used to fit semi-log plots of the data and derive the half-maximal inhibitory concentration (IC_50_) values.

## Author contributions

HMP, PSR, SJ, CGA, and JEC planned experiments. HMP, NK, EA, LH, SD, JR, EB, JD, PSR, SJ performed experiments. GSJ designed SOSV F constructs and provided reagents. CGA, RHC, and JEC supervised research. JEC obtained funding.

## Acknowledgments

We thank Elad Binshtein, Jessica Rodriguez, Chris Gainza, Rachel Sutton, Matthew Goff, Andrew Trivette, and Rachel Nargi of the Vanderbilt Vaccine Center, Paul Rothlauf and Sean Whelan (Washington University in St. Louis), and Tony Schountz and laboratory (Colorado State University) for technical advice and pilot experiments. Support for this work was provided by the Intramural Research Program (National Institute of Allergy and Infectious Diseases). We thank the individual who survived SOSV who participated in the study. The findings and conclusions in this report are those of the authors and do not necessarily represent the official position of the Centers for Disease Control and Prevention.

## Notes

**Conflict of interest statement.** J.E.C. has served as a consultant for Luna Labs USA, Merck Sharp & Dohme Corporation, Emergent Biosolutions, and GlaxoSmithKline, is a member of the Scientific Advisory Board of Meissa Vaccines, a former member of the Scientific Advisory Board of Gigagen (Grifols) and is founder of IDBiologics. The laboratory of J.E.C. received unrelated sponsored research agreements from AstraZeneca, Takeda, and IDBiologics during the conduct of the study. Vanderbilt University has applied for patents for some of the antibodies in this paper.

### Competing Interest Statement

J.E.C. has served as a consultant for Luna Labs USA, Merck Sharp & Dohme Corporation, Emergent Biosolutions, and GlaxoSmithKline, is a member of the Scientific Advisory Board of Meissa Vaccines, a former member of the Scientific Advisory Board of Gigagen (Grifols) and is founder of IDBiologics. The laboratory of J.E.C. received unrelated sponsored research agreements from AstraZeneca, Takeda, and IDBiologics during the conduct of the study. Vanderbilt University has applied for patents for some of the antibodies in this paper.

## REFERENCES

1. Plemper RK, Lamb RA. Paramyxoviridae: the viruses and their replication. In: Howley PM, Knipe DM, Whelan S eds. Fields Virology Volume 1: Emerging Viruses. Kindle Edition: Wolters Kluwer; 2021:503–557

2. Drexler JF, et al. Bats host major mammalian paramyxoviruses. Nat Commun. 2012;3(1):796.

3. Pernet O, et al. Evidence for henipavirus spillover into human populations in Africa. Nat Commun. 2014 51 2014;5(1):1–10.

4. Philbey AW, et al. An apparently new virus (family Paramyxoviridae) infectious for pigs, humans, and fruit bats. Emerg Infect Dis. 1998;4(2):269–71.

5. Murray K, et al. A morbillivirus that caused fatal disease in horses and humans. Science 1995;268(5207):94–97.

6. Yaiw KC, et al. Serological evidence of possible human infection with Tioman virus, a newly described paramyxovirus of bat origin. J Infect Dis. 2007;196(6):884–886.

7. Albariño CG, et al. Novel paramyxovirus associated with severe acute febrile disease, South Sudan and Uganda, 2012. Emerg Infect Dis. 2014;20(2):211–216.

8. Amman BR, et al. A recently discovered pathogenic paramyxovirus, Sosuga virus, is present in Rousettus aegyptiacus fruit bats at multiple locations in Uganda. J Wild. Dis. 2015;51(3):774–779.

9. Ang BSP, Lim TCC, Wang L. Nipah virus infection. J. Clin. Microbiol. 2018;56(6):e01875–17.

10. Marsh GA, Wang L-F. Hendra and Nipah viruses: why are they so deadly?. Curr Opin Virol. 2012;2(3):242–247.

11. Jardetzky TS, Lamb RA. Activation of paramyxovirus membrane fusion and virus entry. Curr Opin Virol. 2014;5:24–33.

12. Aguilar HC, et al. Paramyxovirus glycoproteins and the membrane fusion process. Curr. Clin Microbiol. Rep. 2016;3(3):142–154.

13. Kubota M, et al. Trisaccharide containing α2,3-linked sialic acid is a receptor for mumps virus.. Proc Natl Acad Sci. U. S. A. 2016;113(41):11579–11584.

14. Villar E, Barroso IM. Role of sialic acid-containing molecules in paramyxovirus entry into the host cell: a minireview. Glycoconj J. 2006;23(1–2):5–17.

15. Johnson RI, et al. Characterization of Teviot virus, an Australian bat-borne paramyxovirus. J Gen Virol. 2019;100(3):403–413.

16. Stelfox AJ, Bowden TA. A structure-based rationale for sialic acid independent host-cell entry of Sosuga virus. Proc Natl Acad Sci. U. S. A. 2019;116(43):21514–21520.

17. Kulkarni-Kale U, et al. Mapping antigenic diversity and strain specificity of mumps virus: a bioinformatics approach. Virology. 2007;359(2):436–446.

18. Cusi MG, et al. Localization of a new neutralizing epitope on the mumps virus hemagglutinin–neuraminidase protein. Virus Res. 2001;74(1–2):133–137.

19. Wolinsky JS, Waxham MN, Server AC. Protective effects of glycoprotein-specific monoclonal antibodies on the course of experimental mumps virus meningoencephalitis. J Virol. 1985;53(3):727–34.

20. Kasel JA, et al. Acquisition of serum antibodies to specific viral glycoproteins of parainfluenza virus 3 in children. J Virol. 1984;52(3):828–32.

21. Vaidya SR, Dvivedi GM, Jadhav SM. Cross-neutralization between three mumps viruses & mapping of haemagglutinin-neuraminidase (HN) epitopes. Indian J Med Res.2016;143(1):37–37.

22. Harbury PB, et al. A switch between two-, three-, and four-stranded coiled coils in GCN4 leucine zipper mutants. Science 1993;262(5138):1401–1407.

23. Bose S, et al. Fusion activation through attachment protein stalk domains indicates a conserved core mechanism of paramyxovirus entry into cells. J Virol. 2014;88(8):3925–41.

24. Russell CJ, Jardetzky TS, Lamb RA. Membrane fusion machines of paramyxoviruses: capture of intermediates of fusion. EMBO J. 2001;20(15):4024–34.

25. Bose S, Jardetzky TS, Lamb RA. Timing is everything: fine-tuned molecular machines orchestrate paramyxovirus entry. Virology. 2015;479–480:518–531.

26. Welch SR, et al. Development of a reverse genetics system for Sosuga virus allows rapid screening of antiviral compounds. PLoS Negl Trop Dis. 2018;12(3):e0006326.

27. Van Brussel K, Holmes EC. Zoonotic disease and virome diversity in bats. Curr Opin Virol. 2022;52:192–202.

28. Thibault PA, et al. Zoonotic potential of emerging paramyxoviruses: knowns and unknowns. Adv Virus Res. 2017;98:1.

29. Mousa JJ, et al. Human antibody recognition of antigenic site IV on Pneumovirus fusion proteins. PLOS Pathog. 2018;14(2):e1006837–e1006837.

30. Bowden TR, et al. Menangle virus, a pteropid bat paramyxovirus infectious for pigs and humans, exhibits tropism for secondary lymphoid organs and intestinal epithelium in weaned pigs. J Gen Virol. 2012;93(5):1007–1016.

31. Yaiw KC, et al. Tioman virus, a paramyxovirus of bat origin, causes mild disease in pigs and has a predilection for lymphoid tissues.. Virol. 2008;82(1):565–8.

32. Lan K, et al. Epstein-Barr Virus (EBV): infection, propagation, quantitation, and storage. Curr Protoc Microbiol. 2007;6(1):14E.2.1–14E.2.21.

33. Chen EC, et al. Diverse patterns of antibody variable gene repertoire disruption in patients with amyloid light chain (AL) amyloidosis. PLoS ONE. 2020;15(7 July). doi:10.1371/journal.pone.0235713

34. Schindelin J, et al. Fiji: an open-source platform for biological-image analysis. Nat Methods. 2012;9(7):676–682.

